# Comprehensive 3D Imaging of Whole Zebrafish Using a Water-Based Clearing Reagent for Hard Tissues

**DOI:** 10.1101/2025.04.04.647305

**Authors:** Kiyohito Taimatsu, John Prevedel, Daniel Castranova, Kanako Inoue, Amit Puthan, Brant M. Weinstein

## Abstract

Zebrafish *(Danio rerio)* is a valuable model organism for studying developmental processes due to its external development and the optical clarity of its embryos and larvae. However, as development proceeds zebrafish form increasingly opaque tissues that impede visualization of deep tissues and structures. Although tissue clearing methods have been applied to facilitate imaging at these later stages, most of these methods have limited ability to clear dense tissues such as bone and cartilage, cause significant morphological distortion, and/or result in loss of fluorescent signal when used for imaging of fluorescent transgenes, dye stained animals, or specimens generated using immunofluorescence or fluorescence *in situ* hybridization methods. Here, we report a novel imaging technique using a recently developed clearing reagent called LUCID that makes it possible to capture the complete cellular-resolution 3D structures of larval and juvenile zebrafish. We show that LUCID clears dense tissues such as pharyngeal cartilage in juvenile animals and even tooth bone in adults without causing either significant morphological distortion or significant loss of signal from transgene-driven fluorescent proteins, fluorescent nuclear DNA or actin staining dyes, or whole mount *in situ* hybridization chain reaction (HCR) fluorescence. Using this new approach it is possible to perform complete high-resolution 3D imaging of whole fluorescently stained animals, even deep internal regions, providing a novel tool for elucidating the complex internal structures of developing zebrafish.

## INTRODUCTION

Three-dimensional (3D) imaging and modeling are powerful tools for elucidating the complex morphology of organisms (Rahman et al., 2017, Haleem and Javaid, 2019). However, many model organisms are opaque and grow to sizes that make it difficult to perform whole-body 3D optical imaging and modeling. Zebrafish is widely used in developmental biology due to its external development and small, transparent embryos and larvae (Kimmel et al., 1995), yet capturing the entire body in 3D remains challenging, especially at later stages of development (Kimmel et al., 1998). Organs deep inside the head such as the pharynx become encased in matrix-rich, high-density tissues including cartilage, muscle, and skull bone that cause scattering, reflection, and refraction of incident light, interfering with optical imaging. Non-optical methods such as micro-computed tomography (micro-CT) can be used for whole-body imaging of fixed specimens, but the resolution of this method is much lower than optical imaging and the fixation and staining processes employed often cause notable tissue deformation(Glancy et al., 2023). Furthermore, X-ray–based approaches do not detect fluorescence from fluorescent proteins, stains, or labels, limiting their versatility (Tian et al., 2021).

Tissue clearing technologies have advanced considerably to address these challenges (Tainaka et al., 2016, Tian et al., 2021). One approach to tissue clearing removes or denatures opaque molecules in fixed tissues, such as through defatting or decolorization, while another approach immerses fixed tissue samples in high refractive index (RI) solutions to unify the RI of tissue components, reducing reflection and refraction and enabling deeper light penetration. In practice, these methods are often combined. Indeed, in neuroscience, clearing and fluorescent imaging of thick brain samples are now well established. However, fully reconstructing vertebrate tissues in three dimensions requires clearing bone and cartilage, which remain difficult to clear due to their matrix-rich nature (Utagawa et al., 2023, Jing et al., 2019).The refractive index (RI) of demineralized bone and cartilage is approximately 1.53 (Jing et al., 2019), so a clearing reagent with an RI of at least 1.53 is required to image tissues deep within the heads of older developing animals. Most reagents with RIs this high are organic solvent-based (Jing et al., 2019, Tainaka et al., 2016), causing strong denaturing of proteins, quenching of fluorescent proteins, and distortion of soft tissues. Although water-based clearing reagents induce less protein denaturation, the lower RI of most of these reagents limits the degree of refractive-index matching required for robust clearing of dense matrix-rich tissues. A few relatively high-RI water-based clearing reagents have been reported, including seeDB seeDB1 (1.49) (Ke et al., 2013), seeDB2 (1.52) (Ke et al., 2016), and most recently, LUCID (1.54) (Sekitani et al., 2016, Mizutani et al., 2018, PhotonTech Innovations Co., 2025). However, most water-based reagents underperform organic solvent– based reagents in terms of absolute clarity, and it has been challenging to perform whole-body imaging and full 3D reconstruction of animals once cartilage and bone formation has begun without causing significant degeneration to soft tissues.

In this study we focus on LUCID, a reagent designed specifically to minimize sample distortion that is capable of clearing both soft and hard tissues (Sekitani et al., 2016, Mizutani et al., 2018, PhotonTech Innovations Co., 2025). We show that LUCID clears whole larval and juvenile zebrafish specimens while preserving their overall morphology and the fluorescence resulting from the use of a variety of different modalities. LUCID-cleared zebrafish larvae become almost completely transparent, allowing for detection of all internal cells and structures. GFP and RFP fluorescence in LUCID-prepared transgenic animals can remain stable for over a month, enabling extended and repeated imaging of specimens. Fluorescence dyes such as Hoechst and SiR700-Actin are similarly well-retained. LUCID even enables imaging of entire 24 dpf juvenile zebrafish, and the interior of bone-encased adult teeth. Using LUCID clearing, detailed visualization and three-dimensional modeling of the entire zebrafish body is possible even at these later stages of development, including imaging of previously difficult-to-observe deep bone- and cartilage-encased structures. The “Comprehensive 3D Imaging” methods we have developed for using LUCID together with fluorescent transgenes, dyes, or *in situ* hybridization techniques result in truly whole-body 3-D reconstructions, providing a powerful new tool for furthering our understanding of vertebrate biology.

## METHODS

### Fish husbandry and fish strains

Fish were housed in a large zebrafish-dedicated recirculating aquaculture facility (4 separate 22,000L systems) in 6L and 1.8L tanks. Fry were fed rotifers and adults were fed Gemma Micro 300 (Skretting) once per day. Water quality parameters were routinely measured, and appropriate measures were taken to maintain water quality stability (water quality data available upon request). The following transgenic and mutant zebrafish lines were used for this study: *Tg(mrc1a:eGFP*)*^y251^* (Jung et al., 2017), *Tg(kdrl:mcherry)^y205^*(Fujita et al., 2011), *Tg(fli1a:egfp)^y1^* (Lawson and Weinstein, 2002), and *roy, nacre* (*“casper”*) double mutants (White et al., 2008), crossed into the transgenic backgrounds to eliminate potentially obscuring melanocyte and iridophore cell populations. All zebrafish lines were maintained on the EK wild type background. This study was performed in an AAALAC accredited facility under an active research project overseen by the NICHD ACUC, Animal Study Proposal # 21-015.

### SIR700-Actin Staining

Samples were fixed in 4% paraformaldehyde (PFA) for 16–30 hours. Fixation beyond 36 hours was avoided to reduce autofluorescence. After fixation, the samples were washed twice in 1X PBS for 5 minutes each. Subsequently, they were incubated overnight at 4°C with gentle shaking in 100 nM SIR700-Actin (Cytoskeleton) in PBS. Following staining, the samples were washed twice in 1X PBS for 5 minutes each and then used for observation.

### Nuclear Staining

Samples were fixed in 4% PFA for 16–30 hours. For juvenile samples, the fixative was replaced with fresh 4% PFA approximately 2 hours after immersion to ensure adequate fixation. After fixation, samples were washed twice for 5 minutes each in 1X PBS in the dark on a shaker. They were then stained overnight at 4°C with gentle shaking in 5 µg/mL Hoechst33258 (Latt and Stetten, 1976) or DAPI (Kapuscinski, 1995) in PBS. After staining, samples were washed twice in 1X PBS for 5 minutes each and used for observation. When HCR (see below) was employed, DAPI was mainly used, as permeabilization steps could enhance probe penetration. For samples without HCR treatments, Hoechst was frequently chosen. Lower staining concentrations can result in poor labeling of deep tissue regions, whereas higher concentrations risk unbound dye depositing in tissues and reabsorbing the blue fluorescence signal. Hence, the optimal concentration and volume of Hoechst/DAPI need to be adjusted depending on developmental stage and sample size. We found that DRAQ5 (Smith et al., 2000) was unsuitable for whole-body staining, as the tissue-surface–bound DRAQ5 could absorb the infrared fluorescence from deeper tissue regions (data not shown).

### Hybridization Chain Reaction (HCR) for whole Embryo Staining

HCR was performed to visualize and confirm the identities of each cell type/cluster. HCR probesets were designed by Molecular Instruments (Molecular Instruments, Los Angeles, CA). Embryos were fixed, permeabilized, and hybridized following the steps outlined below.

Approximately 40 embryos were loaded into a 1.6 mL Eppendorf tube containing ice-cold 4% paraformaldehyde (PFA) (Electron Microscopy Sciences Cat #15710) and incubated for 24 hours at 4°C. Following fixation, embryos were washed three times for 5 minutes each with 1 mL of PBST (1X phosphate-buffered saline and 0.1% Tween-20) (Gibco 10010023/Millipore Sigma 9005-64-5). Embryos were then dehydrated through an ascending ethanol series (25%, 50%, 75%, and 96% in PBST), followed by two washes in 100% methanol. Each dehydration step was performed for 15 minutes at 4°C. Embryos were stored at -20°C overnight before further processing.

Embryos were transferred into a 1.6 mL Eppendorf tube and methanol was removed. Permeabilization steps were conducted on a horizontal platform shaker at 20-50 rpm unless otherwise noted. Embryos were washed twice for 5 minutes each in 96% ethanol before incubation in a 1:1 (v/v) ethanol/xylene solution for 1 hour at RT. The samples were subsequently washed twice for 5 minutes each in 96% ethanol, then rehydrated through a descending ethanol series (90%, 75%, 50%, 25%) in PBST for 15 minutes. Embryos were washed twice for 5 minutes each in PBST.

For permeabilization, embryos were incubated in 80% acetone at -20°C for 30 minutes without shaking. Embryos were washed twice for 5 minutes each in PBST and bleached in 3% H₂O₂ / 1% KOH in PBST for 15-45 minutes, monitoring for sufficient depigmentation. Embryos were washed in PBST for 5 minutes, followed by an additional 10-minute wash.

Fixed embryos were pre-hybridized with 500 μL of preheated (37°C) HCR probe hybridization buffer (Molecular Instruments, HCR™ RNA-FISH Bundle) for 30 minutes at 37°C. Probe hybridization was performed by diluting 2 pmol of each 1 μM probe set in 500 μL of hybridization buffer, replacing the pre-hybridization buffer, and incubating the embryos at 37°C for 12-16 hours. Excess probes were removed by washing embryos four times for 15 minutes each in preheated HCR probe wash buffer (Molecular Instruments, HCR™ RNA-FISH Bundle) at 37°C. This was followed by two 5-minute washes with 5X SSCT (12.5 mL of 20X SSC in 50 μL Tween-20) (KD Medical Cat # RGF-3240 / Millipore Sigma 9005-64-5) at RT.

Embryos were pre-amplified with 500 μL of HCR probe amplification buffer (Molecular Instruments, HCR™ RNA-FISH Bundle) for 30 minutes at RT. Hairpins were prepared separately by diluting 12 pmol of hairpin h1 and hairpin h2 (4 μL of 3 μM stock each) in PCR tubes. Hairpins were snap-cooled by heating at 95°C for 90 seconds, followed by cooling to RT in the dark for 30 minutes. The snap-cooled hairpins were then mixed in 200 μL of fresh HCR probe amplification buffer at RT. The pre-amplification buffer was removed, and the hairpin solution was added to the embryos. Hybridization with hairpins was conducted in the dark at RT for 12-16 hours.

Excess hairpins were removed by washing embryos with 500 μL of 5X SSCT at RT in the following sequence: two 5-minute washes, two 30-minute washes, and a final 5-minute wash. At this stage, samples were optionally stained with 50 ug/ml DAPI or Hoechst. Embryos were stored at 4°C. Then, the samples were washed twice with PBS for 5 minutes each.

### Sample clearing with LUCID

For the clearing of fixed embryo samples in PBS, PBS was removed from the tube, followed by the careful removal of residual moisture from the bottom of the tube using a twisted Kimwipe (Kimberly-Clark). The embryos were then incubated in 125 μL of 100% LUCID solution (PhotonTech Innovations Co) per tube for 24 hours at 4°C, protected from light.

For the clearing of fixed juvenile samples in PBS, decalcification was done followed using 0.5 M EDTA pH 8.0 (Invitrogen) at 4°C for 2 days on a shaker (Jing et al., 2018, Jing et al., 2019). Next, 25% v/v Quadorol (Sigma-Aldrich 360538) and 5% v/v ammonium were used for decolorization at 4°C for 2–4 days on a shaker to remove hemoglobin (Tainaka et al., 2018, Jing et al., 2018). Defatting was then performed in a graded series of 25%, 50%, 70% v/v tert-BtOH (Sigma-Aldrich 360538), each containing 3% w/v Quadorol, at room temperature for 24 hours on a shaker (Jing et al., 2018). Finally, samples were transferred back to PBS.

The samples were removed, and excess moisture was gently wiped off using a Kimwipe. The samples were transferred to 25% LUCID solution and sequentially replaced with 50% and 75% LUCID solutions at 4°C, with each step performed for 1 hour on a shaker. Subsequently, the samples were transferred to 100% LUCID solution and incubated at 4°C on a shaker for 12 hours. The solution was then replaced with fresh 500 μL of 100% LUCID, and the samples were incubated at 4°C on a shaker for an additional 24 hours to complete the clearing process.

### Mounting

Samples were mounted in LUCID for imaging. Samples were placed in a plastic dish containing LUCID, and pigmented eyes were removed using forceps or a tungsten needle. The removal of eyes was more challenging prior to LUCID treatment. A glass-bottom dish was prepared, and a well was created using a double layer of vinyl tape. The sample was placed within the well, and a coverslip cleaned with 70% ethanol was placed over the well. The edges of the coverslip were sealed with nail polish to prevent LUCID from evaporating.

### Imaging

Samples were imaged using a Nikon Ti2 inverted microscope equipped with a Yokogawa CSU-W1 spinning disk confocal (Hamamatsu Orca Fusion-BT camera). The laser output was set to 25%. Four channels were used for fluorescence detection: Blue (405 nm) for autofluorescence in matrix-rich tissues or Hoechst/DAPI, Green (488 nm) for GFP, Red (561 nm) for RFP, RBC autofluorescence, or HCR probe signals, and Infrared (640 nm) for gut autofluorescence or other dyes/HCR probe signals. The exposure time was 80 ms, and the Z-step size was 1 µm for a 10× lens, 0.6 µm for a 20× lens, and 0.4 µm for a 20× lens. A 10× air objective lens (NA 0.45) was mainly used for imaging entire embryos or juveniles, a 20× water-immersion objective lens (NA 0.95 or 1.0) was employed for detailed imaging of embryonic heads, while a 40× water-immersion objective lens (NA 1.15) was employed for nuclear level imaging of embryonic heads. Some samples were observed using a stereomicroscope (Leica M165, Leica DFC 7000T camera, Leica Application Suite X software), with images were acquired in either 12-bit or 16-bit format.

### Image Processing

Image processing was primarily conducted using NIS Elements. First, tile-scanned images were stitched with stitching software (NIS Elements), and images were flipped as needed to match sample orientation. AI-based noise reduction was then applied using Nikon Denoise AI. Focus stacking using Nikon Elements’ Extended Depth of Field was performed on some transmitted light images.

Three-dimensional models were also created during image processing. For images with a large dynamic range, non-linear adjustments (gamma correction) were sometimes applied to enhance visibility. The 3D models were oriented appropriately in three-dimensional space, and necessary images were captured. Half-body images and optical sections were generated by removing the contralateral half of the 3D model or by excluding undesired optical sections, respectively. The thickness of the optical sections was generally equivalent to 3–5 cell layers, depending on the region of interest. Movies were recorded by rotating the 3D model in NIS Elements. Sequences were spliced and labels were added to movies using Adobe Premiere Pro 2024.

## RESULTS

Zebrafish embryos and larvae are widely used for high-resolution imaging of developmental processes because of their optical clarity. However, as development proceeds imaging of deep tissues becomes progressively hampered by light scattering and by the presence of opaque tissues such as bone and cartilage. As noted above, clearing reagents have been used to aid visualization of deeper structures in older, more opaque animals, but most of these agents have limited ability to clear dense tissues such as bone and cartilage, cause significant morphological distortion, and/or result in loss of fluorescent signal when used for imaging of transgenic fluorescent proteins, fluorescent dye stained animals, or specimens stained using immunofluorescence or fluorescence *in situ* hybridization methods. We have tested the ability of a new agent, LUCID (Sekitani et al., 2016, PhotonTech Innovations Co., 2025), to clear dense tissues without introducing significant morphological distortion or causing loss of fluorescence generated using a variety of modalities.

Uncleared five day post-fertilization (5 dpf) zebrafish larvae are largely translucent for purposes of imaging the trunk, especially when using pigment-deficient *casper* animals (**Fig. 1A**), but cartilage and other opaque tissues have already begun to interfere with imaging of deeper tissues in the head by 5 dpf (**Fig. 1C**). This is even more evident in confocal images of networks of blood and lymphatic vessels, readily visualized using *casper* mutant, *Tg(kdrl:mcherry)^y206^*, *Tg(mrc1a:egfp)^y251^* double-transgenic reporter lines (Fujita et al., 2011, Jung et al., 2017) that label blood (magenta) and lymphatic (green) endothelium, respectively (**Fig. 1E,G**). Cranial vessels closest to the objective can be imaged nicely in 5 dpf animals, but resolution fades quickly and all but disappears in the half of the head furthest from the objective (**Fig. 1E,G**). In contrast, LUCID-cleared 5 dpf zebrafish larvae are far more transparent, appearing almost invisible to the eye (**Fig. 1B**), even when imaging through their heads (**Fig. 1D**). Confocal imaging of LUCID-cleared 5 dpf *casper, Tg(kdrl:mcherry)^y206^*, *Tg(mrc1a:egfp)^y251^*double-transgenic animals shows that mCherry and EGFP signals are both very well-preserved (**Fig. 1F,H**, **Movie 1**), and that LUCID clearing permits effective imaging through the entire depth of the head (**Fig. 1F,H**, **Movie 1**), with complete reconstruction of both the blood and lymphatic vasculature (**Fig. 1F,H**, **Movie 1**). Vessels on the far side of head are imaged with nearly as much clarity and fluorescence signal as vessels on the near side of the head. Thus, applying LUCID to zebrafish embryos enables Comprehensive 3D Imaging while minimizing tissue distortion.

**Figure 1.**
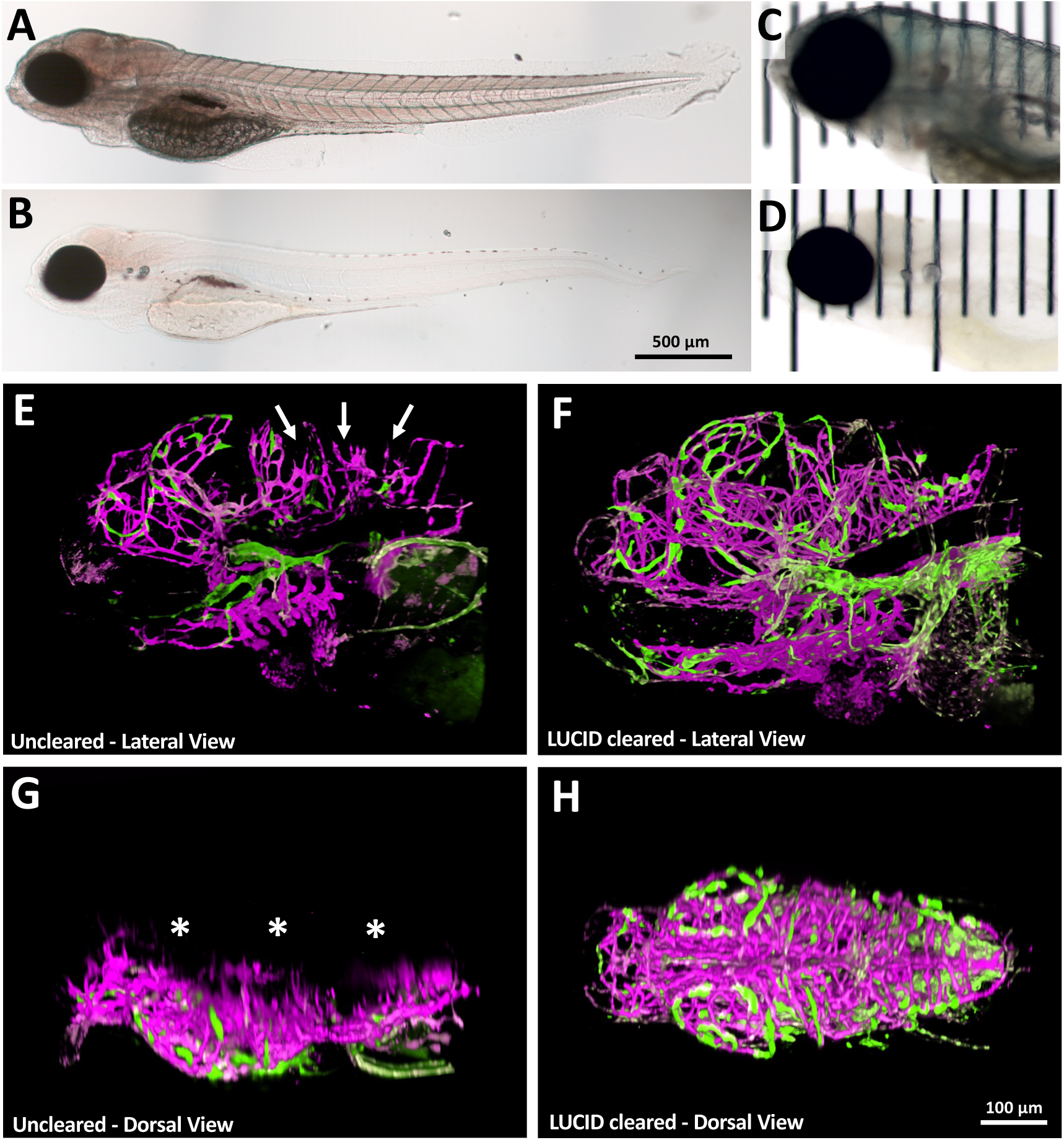
LUCID clearing facilitates deep imaging of zebrafish larvae. **A,B,** Transmitted light images of whole uncleared (A) or LUCID cleared (B) 5 dpf larvae. **C,D,** Higher magnification transmitted light images of the head and rostral trunk of uncleared (C) or LUCID cleared (D) 5 dpf larvae. Animals are photographed on top of a micrometer to allow for assessment of their clarity. **E-H,** Reconstructions of confocal image stacks collected from the left lateral side of the heads of uncleared (E,G) or LUCID cleared (F,H) 5 dpf *casper* mutant, *Tg(kdrl:mcherry)^y206^*, *Tg(mrc1a:egfp)^y251^* double-transgenic larvae, showing mCherry (magenta) and EGFP (green) fluorescence. The heavily pigmented eyes were removed to permit imaging through the head. Panels E and F show lateral reconstructions of the heads, while panels G and H show dorsal views generated from the same image stacks. In comparison to LUCID cleared animals far fewer hindbrain vessels are visible in the uncleared animals (arrows in E) and vessels are not seen at all (asterisks) on the opposite (right) side of the head from the side that the images were collected from (the left side). Scale bars = 500 µm A,B) and 100 µm (E-H). For E-H, also see **Supplemental Movies 1,2**.

To expand the utility of Comprehensive 3D Imaging, a method for uniformly staining tissues— including deeper regions—is essential. Traditional histological stains such as H&E or Masson’s trichrome are often unsuitable because they can obscure deeper layers once surface staining is achieved. By contrast, fluorescent labeling allows the sample to remain transparent overall, and only the targeted molecules are detected when excited by the laser. As a representative fluorescent stain we performed whole-body nuclear staining with DAPI (F**ig. 2A–D**). A single optical slice near the midline yields details that resemble a resin-embedded section stained with DAPI, yet here it is part of a complete 3D dataset with minimal sample damage. We typically use 5 µg/mL DAPI or Hoechst. Lower concentrations result in poor labeling of deep tissues, whereas higher concentrations can lead to non-specific deposition of dye that reabsorbs blue fluorescence (**Fig. 2E–H**). The concentration must be balanced for each developmental stage and sample size. We also found DRAQ5 to be unsuitable for whole-body staining because surface-bound DRAQ5 absorbs infrared fluorescence from deeper tissue regions. We also tested the near-infrared fluorescent actin probe SiR700-Actin (Cytoskeleton). LUCID clearing preserved SiR700-Actin signals in jaw, pharynx, and trunk muscles across the entire sample (**Fig. 2E–H**, **Movie 2**).

**Figure 2.**
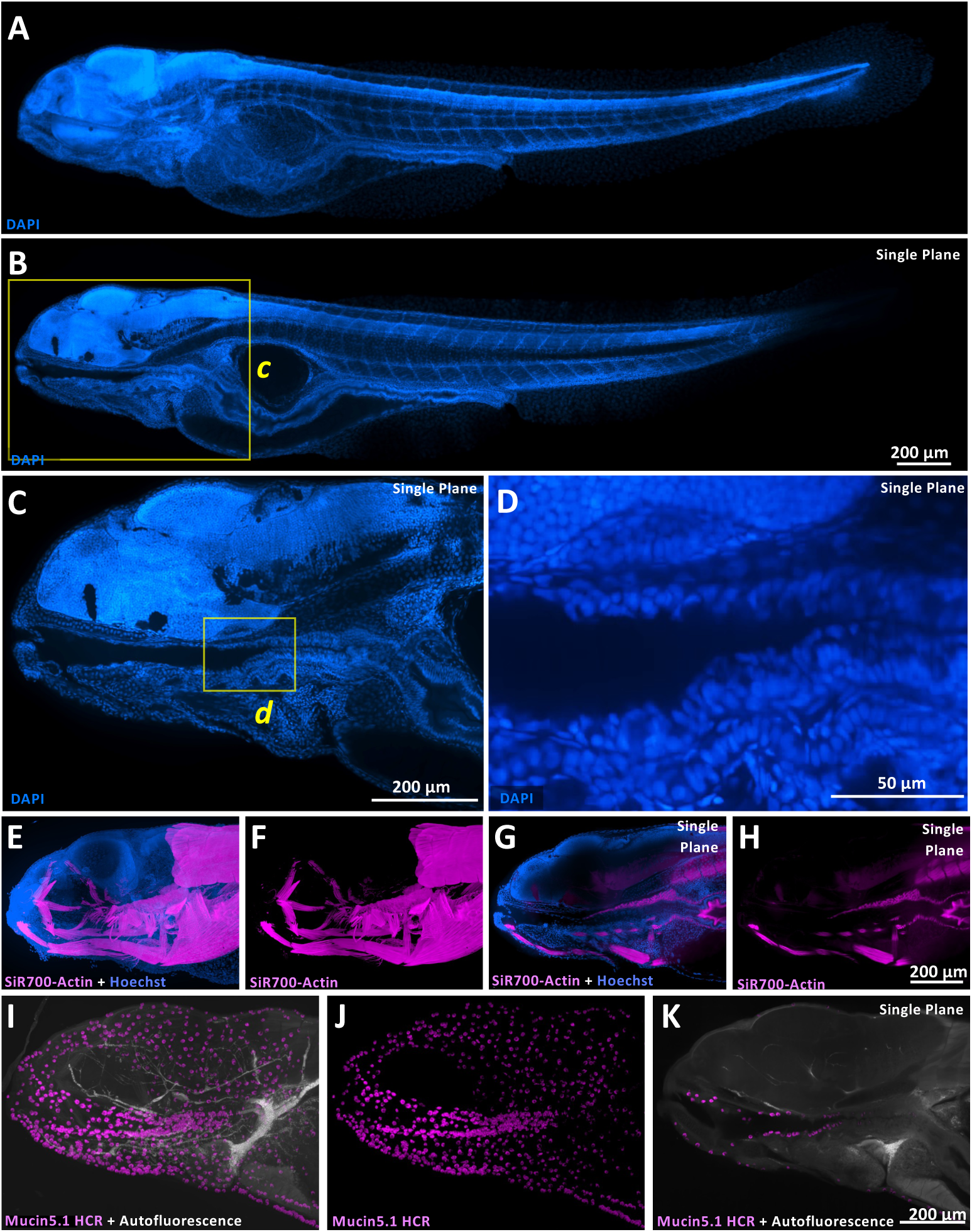
Whole body fluorescent staining and comprehensive 3D imaging. **A-D,** Confocal images of a LUCID-cleared 5 dpf DAPI-stained *casper* zebrafish larva, showing (A) an alpha blending projection of the entire image stack, (B) a single confocal image plane from the center of the stack at the level of the pharynx, (C) a magnified portion of the confocal image plane from the boxed region in panel B, and (D) a magnified portion of the confocal image plane from the boxed region in panel C, highlighting clear cellular-level resolution of the deep, cartilage-enclosed pharynx. **E-H,** Confocal images of a LUCID-cleared 5 dpf Hoechst 33342- and SiR700-Actin-stained *casper* zebrafish larva, showing alpha blending projection of the entire image stack with (E) Hoechst 33342 and SiR700-Actin together or (F) SiR700-Actin alone, or a single confocal image plane from the center of the stack approximately at the level of the pharynx with (G) Hoechst 33342 and SiR700-Actin together or (H) SiR700-Actin alone. Actin-rich muscle structures are clearly visualized in the center of the head. **I-K,** Confocal images of a LUCID-cleared 5 dpf *casper* zebrafish larva subjected to whole mount *in situ* hybridization chain reaction (HCR) probing for *mucin5.1*, showing alpha blending projection of the entire image stack with (I) both *mucin5.1* HCR fluorescence (magenta) and erythrocyte autofluorescence (grey) highlighting the vasculature, or only (J) *mucin5.1* HCR fluorescence (magenta), or (K) a single confocal image plane from the center of the stack at the level of the pharynx including both *mucin5.1* HCR fluorescence (magenta) and erythrocyte autofluorescence (grey). *Mucin5.1* HCR-positive goblet cells lining the pharynx are clearly visualized. The heavily pigmented eyes were removed from all animals to permit imaging through the head. All images are lateral views, rostral to the left. Image reconstructions are generated from “stitched” stacks combined from separate confocal stacks collected from overlapping adjacent areas of the same animal. Scale bars = 200 µm (A,B,C,E-K) and 50 µm (D

Beyond detecting cellular structures, we sought a method to visualize gene expression patterns throughout the entire sample. Hybridization chain reaction (HCR) is a recently popularized fluorescent *in situ* hybridization approach that offers simpler workflows than traditional whole-mount in situ hybridization (WISH), quantitative sensitivity, and the potential for multiplexing (Choi et al., 2010). Because it typically involves less heat denaturation, HCR is well suited for analyzing developmental processes and complex tissue architectures. A major obstacle is achieving uniform probe penetration into deeper regions without damaging the sample surface— especially in older embryos. By incorporating a xylene/acetone protocol to enhance tissue permeability (Vauti et al., 2020), we successfully labeled transcripts in the center of zebrafish embryos (**Fig. 2I–K**, **Movie 3**). For example, mucin5.1 expression in pharyngeal goblet cells at 5 dpf was clearly visualized, revealing a continuous distribution of secretory cells from the external epithelium to the deeper pharyngeal area.

LUCID provides even greater advantages in older animals. Improved deeper imaging of 5 dpf animals is useful, but imaging at depth becomes far more challenging in juvenile and adult fish. After LUCID-clearing a 24 dpf, Hoechst 33342 dye-stained juvenile zebrafish we collected 655 lateral-view confocal image sections through the entire width of the animal, imaging through a depth of 655 microns (**Fig. 3A**, **Movie 4**). The complete 3D structure of the juvenile fish was visualized without apparent damage or shrinkage, including deep internal structures, capturing both the overall body plan and detailed histology-like sections of various internal structures (**Movie 4**). By 24 dpf the morphology of the developing zebrafish resembles that of the adult fish, with many adult-like organs and tissues. An optical section roughly through the trunk midline (**Fig. 3B**) provides clear visualization of the structure of the rhombomeres of the hindbrain (**Fig. 3C**), the swim bladder and its epithelial layers and the pronephric duct (**Fig. 3D**), and the heart chambers including the ventricle and bulbus arteriosus (outflow tract) (**Fig. 3E**). Structures that traverse multiple optical sections can also be readily visualized and traced through these optical sections and visualized in 2D by generating composites or “collages” of several aligned adjacent optical sections, as shown for the pneumatic duct, a structure used by fish to introduce air into the swim bladder in order to maintain neutral buoyancy (**Fig. 3F**). We can clearly visualize and continuously trace the thin pneumatic duct opening all the way from its starting point in the zebrafish pharynx to its terminus in the swim bladder (asterisks in **Fig. 3F**). We are also able to image the developing zebrafish gonads (**Fig. 3G,H**) and their immature oocytes (**Fig. 3H**, arrowhead). Other deep structures are also similarly well visualized (**Fig. 3I-L**), such as developing gill filaments in the ventral jaw (**Fig. 3J**), the area of the anal pore with separate but closely juxtaposed openings from the intestine, kidneys, and gonads (**Fig. 3K**), and the developing brush border in the wall of the intestine (**Fig. 3L**). Each optical slice reveals precise morphology and positioning akin to a histological section, yet these are optical sections covering the entire body via tomographic scanning, avoiding tissue damage or distortion associated with histological processing and mechanical sectioning and allowing for a full reconstruction of the entire animal.

**Figure 3.**
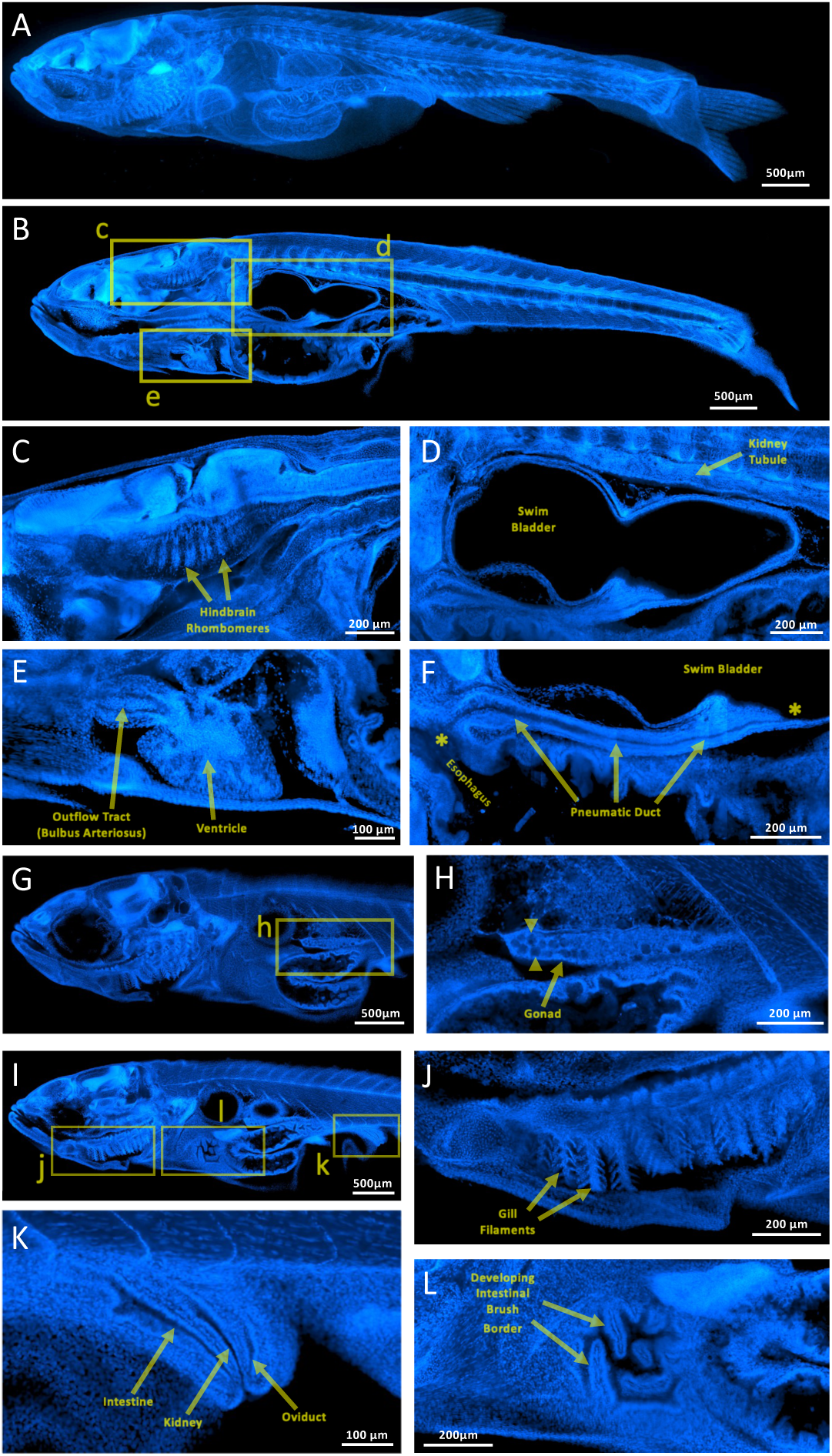
LUCID clearing facilitates deep imaging of juvenile zebrafish. Confocal image reconstructions of a LUCID-cleared, Hoechst 33342-stained 24 dpf juvenile zebrafish. Image reconstructions are generated from a single “stitched” stack combined from 12 separate confocal stacks collected from overlapping adjacent areas of the same animal, with 655 planes per stack each separated by a Z-distance of 1 µm for a total depth of 655 µm, and with total X-Y dimensions of 13762 x 3199 pixels (8,945 µm x 2,079 µm). **A,** Maximum intensity projection of the entire stitched image stack. **B,** Single confocal image plane from the center of the stack at the level of the pharynx (at a depth of 335 µm in the stack), with boxes noting magnified areas shown in panels C-E. **C-E,** Magnified areas of the confocal image plane in panel B, showing sections through the hindbrain rhombomeres (C), swim bladder and kidney tubule (D), and heart ventricle and bulbus arteriosus (E). **F,** Collage of 10 magnified sections from the region just below the swim bladder collected from 287µm to 350 µm deep in the image stack, showing the complete pneumatic duct linking the esophagus to the swim bladder (openings to the esophagus and swim bladder noted with asterisks). **G,** Single confocal image plane from the rostral stack at the level of the left gonad (at a depth of 198 µm in the stack), with a boxes noting the area magnified in panel H. **H,** Magnified area of the confocal image plane in panel G, showing the developing gonad with immature oocytes (arrowheads). **I,** Single confocal image plane from the stitched stack at the level of left-mid side (at a depth of 225 µm in the stack), with boxes noting magnified areas shown in panels J-L. **J-L,** Magnified areas of the confocal image plane in panel I, showing sections through gill filaments (J), the anal pore (K), and the developing brush border in the wall of the intestine (L). All images are lateral views, rostral to the left. Scale bars = 500 µm (A,B,G,I), 200 µm (C,D,F,H,J,L), and 100 µm (E,K).

Finally, to demonstrate LUCID’s capability to clear hard and soft tissues simultaneously, we imaged zebrafish teeth along with surrounding tissues. Although other currently available reagents have the capacity to clear soft tissue, bone and other hard tissues remain extremely challenging, especially bony adult tissues such as teeth. Zebrafish possess pharyngeal teeth with an anatomical structure and cellular composition resembling that of mammalian teeth, but unlike mammals zebrafish replace their teeth approximately every two weeks throughout adult life (Wautier and Huysseune, 2001), making zebrafish an excellent model for studying tooth formation (**Fig. 4A**). Like the teeth of mammals, zebrafish teeth are highly opaque, obscuring imaging of the tooth interior (**Fig. 4B**), but LUCID clearing of dissected adult lower jaws with their associated teeth renders teeth nearly transparent (**Fig. 4C**). Confocal imaging of uncleared jaws of adult *Tg(fli1a:egfp)^y1^*transgenic animals with EGFP “tagged” vascular endothelial cells permits visualization of more superficial vessels in the jaw (**Fig. 4D,F**), but does not allow clear imaging of vessels in the interior of the tooth, particular vessels deep near the root of the tooth (large arrows in **Fig. 4D,F**). In contrast, distal tooth vessels (small arrows in **Fig. 4E,G**) and vessels deep in the root of the tooth (large arrows in **Fig. 4E,G**) are both readily visualized in high detail and resolution in the interior of LUCID-cleared teeth from *Tg(fli1a:egfp)^y1^* transgenic animals.

**Figure 4.**
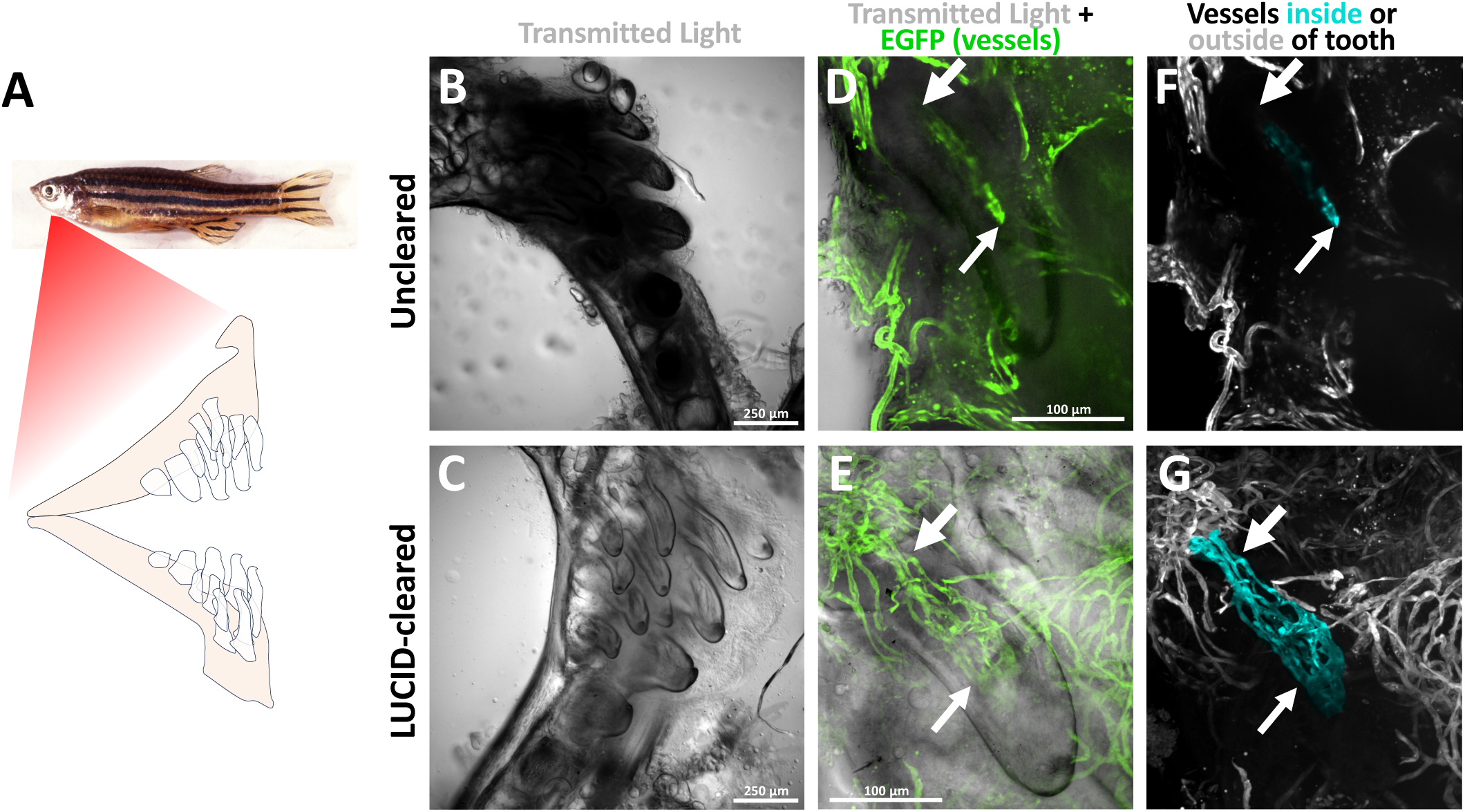
LUCID clearing facilitates imaging through adult teeth. **A,** Image of a fish showing the approximate location of the jaw and pharyngeal teeth (shown in schematic view). **B-G,** Transmitted light focus stacked (B,D,C,E) and/or green confocal vascular (D,F,E,G) imaging of jaws and teeth from uncleared (B,D,F) or LUCID-cleared (C,E,G) adult *Tg(fli1a:egfp)^y1^* transgenic zebrafish. Small arrows note vessels near tips of teeth, large arrows point to regions of deep vessels near base of teeth, readily visualized in LUCID-cleared teeth (panels E,G) but obscured in uncleared teeth (panels D,F). Scale bars = 250 µm (B,C) and 100 µm (D-G).

## DISCUSSION

Our proposed “Comprehensive 3D Imaging” concept goes beyond conventional deep imaging by enabling truly organism-wide 3D visualization and modeling. As a practical realization of this concept, we describe a usage of LUCID for larval and juvenile zebrafish that enables deep 3D imaging and modeling of the entire organism, with excellent preservation of morphology and retention of fluorescence from transgenic fluorescent proteins, fluorescent dyes, and fluorescent *in situ* hybridization. The method allows imaging of whole juvenile animals with tomographic sectioning like X-ray based microCT imaging, but with much higher resolution, less sample damage, and with the ability to visualize fluorescence. Optical sections from Hoechst dye-stained cleared animals provide resolution akin to histological sections, but they are obtained by serial optical sectioning without introducing physical tissue damage or distortion, permitting detailed 3-D reconstruction of tissues and organs. In addition to imaging whole larval and juvenile fish, we show that LUCID clearing can also be used to image the interior of normally opaque dissected adult structures such as teeth, without the necessity of physical sectioning.

Our findings suggest that LUCID clearing is likely to be compatible with most commonly used methods for fluorescence imaging of tissues. Since EGFP and mCherry proteins are both retained in LUCID cleared tissues, we believe that the same will hold true for other fluorescent proteins, especially those derived from EGFP, although this needs to be confirmed by additional imaging of transgenic lines expressing additional fluorescent proteins. Similarly, Hoechst 33342 and SiR700-Actin dyes are retained after LUCID treatment, as is DAPI (unpublished findings), suggesting LUCID treatment does not quench or remove labeling by commonly used fluorescent tissue dyes. Fluorescence from whole mount *in situ* hybridization via HCR is also retained using multiple different secondary fluorescent hairpins (unpublished findings), again suggesting fluorescence retention is not restricted to one fluorescent moiety or channel. Again, however, further testing will be needed to determine whether LUCID clearing is indeed effective when combined with the full spectrum of fluorescent tools used for visualization of fixed tissue specimens.

Optical imaging at depth has intrinsic limitations that even clearing agents can in the end not overcome. Although we have been able to image larval zebrafish up to 24 dpf, optical imaging beyond this stage remains challenging due to the difficulties of such deep imaging even in these highly transparent, refractive index-matched samples. In terms of zebrafish body length, 10 mm is likely the size limit for comprehensive 3D imaging using a spinning disk confocal microscope. The application of additional microscopic imaging methods such as orthogonal imaging via Selective Plane Illumination Microscopy (SPIM) might allow for effective imaging of still older and larger LUCID-cleared specimens (Mizutani et al., 2018) (PhotonTech Innovations Co., 2025), but pushing the limits of tissue clearing to still more advanced stages of development may require new clearing reagents that offer even higher refractive indices while maintaining low protein denaturation.

We performed imaging in this study using *casper* (*roy, nic* double mutant) animals lacking melanocytes and iridophores (White et al., 2008), although these animals still retain melanin-containing retinal pigment epithelium. We removed the eyes to perform imaging of structures behind them, but *slc45a2 (albino)* or other mutants could be used to eliminate melanin pigment without having to remove the eyes, although this would require crossing these mutants into the background of whatever transgenic lines are being used for visualization. Phenylthiourea (PTU) treatment of developing animals has been used as an alternative method to suppress melanin pigment formation, but it is associated with developmental abnormalities that become much more severe and eventually lethal as larval development proceeds (Choi et al., 2007, Li et al., 2012). Hydrogen peroxide treatment of fixed samples is also sometimes used to reduce pigmentation in stained samples, but it can severely damage tissues and frequently quenches fluorescent proteins, so its use is not advisable if one aims to avoid sample deterioration (Costa et al., 2019). In lieu of laborious crossing of transgenic lines onto pigment mutant backgrounds, injection of CRISPR duplex guide ribonucleoproteins targeting the *slc45a2 (albino)* locus has been shown to be an effective means for generating melanin-deficient animals for microscopic imaging (Davis et al., 2021).

In conclusion, we introduce a Comprehensive 3D Imaging framework that achieves high-resolution, whole-body 3D imaging of zebrafish embryos and juveniles, including bone and cartilage. We also establish multiple fluorescent labeling protocols for uniformly staining these samples. Wide application of this method together with registration of the resulting image data sets could be used to generate “atlases” revealing the detailed, correlated 3D distribution of RNAs, proteins, and cell types from embryonic through juvenile stages of development, surpassing the limitations of conventional micro-CT. The method can also be applied to many other organisms and tissue samples, broadening its potential impact. We anticipate that wider use of this method will contribute not only to zebrafish studies but also to the broader field of intact tissue imaging and 3D atlas construction.

## SUPPLEMENTAL MOVIE LEGENDS

**Supp. Movie 1**

Movie showing 3-D reconstruction image views of stitched confocal image stacks of the heads and rostral trunks of uncleared (left) and LUCID-cleared (right) 5 dpf *casper* mutant, *Tg(kdrl:mcherry)^y206^*, *Tg(mrc1a:egfp)^y251^* double-transgenic larvae (images collected from the left side). Blood vessels are in magenta and lymphatic vessels are in green. Images correspond to those shown in Fig. 1E**-H**. Note failure to image vessels on the far (right) side in uncleared animals compared to LUCID-cleared animals, where these vessels are readily visualized.

**Supp. Movie 2**

Movie showing 3-D reconstruction image views of a confocal image stack of the head and rostral trunk of LUCID-cleared 5 dpf Hoechst 33342- and SiR700-Actin-stained zebrafish larva, with nuclei in blue and muscle and other actin-rich tissues in magenta. Images correspond to those shown in **Fig. 2E-H**.

**Supp. Movie 3**

Movie showing 3-D reconstruction image views of a confocal image stack of the head and rostral trunk of a LUCID-cleared 5 dpf zebrafish larva subjected to whole mount *in situ* hybridization chain reaction probing for *mucin5.1*. with erythrocyte autofluorescence in vessels in white and goblet cell mucin5.1 expression in magenta. Images correspond to those shown in **Fig. 2I-K**.

**Supp. Movie 4**

Movie showing 3-D reconstruction image views of the stitched, combined confocal image stack of a LUCID-cleared 24 dpf Hoechst 33342-stained juvenile zebrafish. Images correspond to those shown in **Fig. 3**.

